# Comprehensive cellular analysis with single-nucleus RNA-seq of archived PAXgene whole blood samples

**DOI:** 10.1101/2025.03.06.641241

**Authors:** Ojasvi Chaudhary, Mia Steinberg, Grant Duclos, Peter Gathungu, Manisha Rao, Rogelio Aguilar, Varsha Shankarappa, Chris Rands, Xi Chen, Rebecca Halpin, Elizabeth Galery, Joseph Boland, Maurizio Scaltriti, Brian Dougherty, Asaf Rotem

## Abstract

No method has been developed for single-cell analysis of the large repositories of preserved whole blood samples stored in PAXgene Blood RNA tubes. To address this gap, two nuclei isolation techniques for single-nucleus RNA sequencing were evaluated: mechanical separation (MS), using an Acrodisc filter, and cell lysis (CL). While both methods captured nuclei from all major immune cell types, CL resulted in up to two orders of magnitude higher nuclei yields and less biased proportions of immune cells than MS. High ambient globin gene counts following CL were substantially reduced by CRISPR-guided globin gene depletion of complementary DNA, resulting in more sensitive and efficient gene detection per cell. Despite capturing only nuclear transcripts, CL-derived samples maintained similar cell-type proportions and gene expression as matched peripheral blood mononuclear cell samples, while retaining granulocytes. The CL isolation with globin depletion method enables comprehensive analysis of PAXgene whole blood samples at single-cell resolution.

**MOTIVATION:** PAXgene Blood RNA Tubes are commonly used for blood preservation and gene expression studies due to the ease of use and stabilization of RNA. To date, transcriptomic studies using PAXgene tubes have been limited to bulk RNA profiling approaches such as RNA sequencing (RNA-seq). Here we introduce a method for recovering high-quality nuclei from frozen blood preserved in PAXgene tubes for single-nucleus RNA-seq (snRNA-seq). Using this approach, all white blood cell types – including granulocytes – are captured, thereby enabling comprehensive transcriptomic profiling of peripheral blood at single-cell resolution.

## INTRODUCTION

Longitudinal blood sampling is a minimally invasive method for monitoring patients in clinical settings. Peripheral blood serves as an indicator of immune response and can be used as an early indicator of disease^1^ or as a predictor of treatment response.^2^ Traditionally, peripheral blood mononuclear cells (PBMCs) are isolated from whole blood for subsequent transcriptomic analysis.^3^ However, PBMC analysis has inherent limitations, such as exclusion of granulocytes, which comprise 50–70% of white blood cells (WBCs), the presence of red blood cells (RBCs), and potential RNA degradation or altered gene expression due to processing,^4^ freezing times, storage, and shipment conditions (Table 1). Additionally, PBMCs must be isolated from fresh blood before freezing, which is not always feasible at clinical sites. Often, whole blood is collected using PAXgene Blood RNA Tubes, which contain proprietary reagents to stabilize RNA, offer easy processing at collection sites with minimal training, and can be stored at room temperature for up to 3 days before long-term frozen storage.^5^

**Table 1.** Comparison of methodologies for scRNA-seq of PBMCs and snRNA-seq of nuclei isolated from frozen PAXgene whole blood samples.

|  | scRNA-seq of PBMCs | snRNA-seq of preserved blood |
| --- | --- | --- |
| **Collection** | Indirect, complex, and time consuming | Direct, simple, and quick |
| **Storage** | Immediately at $-80^{\circ}\text{C}$ & $-196^{\circ}\text{C}$ | Flexible; 3 days at RT, 5 days at $4^{\circ}\text{C}$, and long-term at $-20^{\circ}\text{C}$ and $-80^{\circ}\text{C}$ |
| **Stability** | Sensitive to processing time | Stable |
| **Isolate** | Cells | Nuclei |
| **Cell types** | Subset of immune cell | All immune cell types |
| **Single-cell assay limitations** | None | Protein expression and ATAC-seq not tested |

ATAC-seq, assay for transposase-accessible chromatin using sequencing; PBMC, peripheral blood mononuclear cell; RT, room temperature; scRNA-seq, single-cell RNA sequencing; snRNA-seq, single-nucleus RNA sequencing.

The high integrity of RNA in PAXgene tubes enables advanced analysis at single-cell resolution, which can reveal insights into disease biomarkers, rare cell types, cell composition, developmental pathways, cell lineage tracing, and gene regulation.^6,7^ To our knowledge, single-nucleus RNA sequencing (snRNA-seq) has not been performed on frozen PAXgene whole blood samples. This paper presents a novel method for isolating nuclei from frozen PAXgene RNA Blood Tubes for subsequent snRNA-seq analysis. This procedure represents a novel approach to analyze all immune cell types, including granulocytes, from frozen blood samples.

## RESULTS

### Evaluation of frozen PAXgene whole blood samples

Whole blood samples were obtained from three healthy individuals (Donors A, B, and C) and aliquoted for both PBMC isolation and PAXgene processing before being frozen (Figure 1). Analysis of the PAXgene tubes revealed that the freeze–thaw cycle combined with the PAXgene RNA stabilization reagents compromised cell membranes, as indicated by the lack of green fluorescence from acridine orange dye on confocal microscopy imaging of blood pellets from PAXgene whole blood samples resuspended in buffer (Figure S1A). Nuclei remained intact and suitable for snRNA-seq.

**Figure 1.**
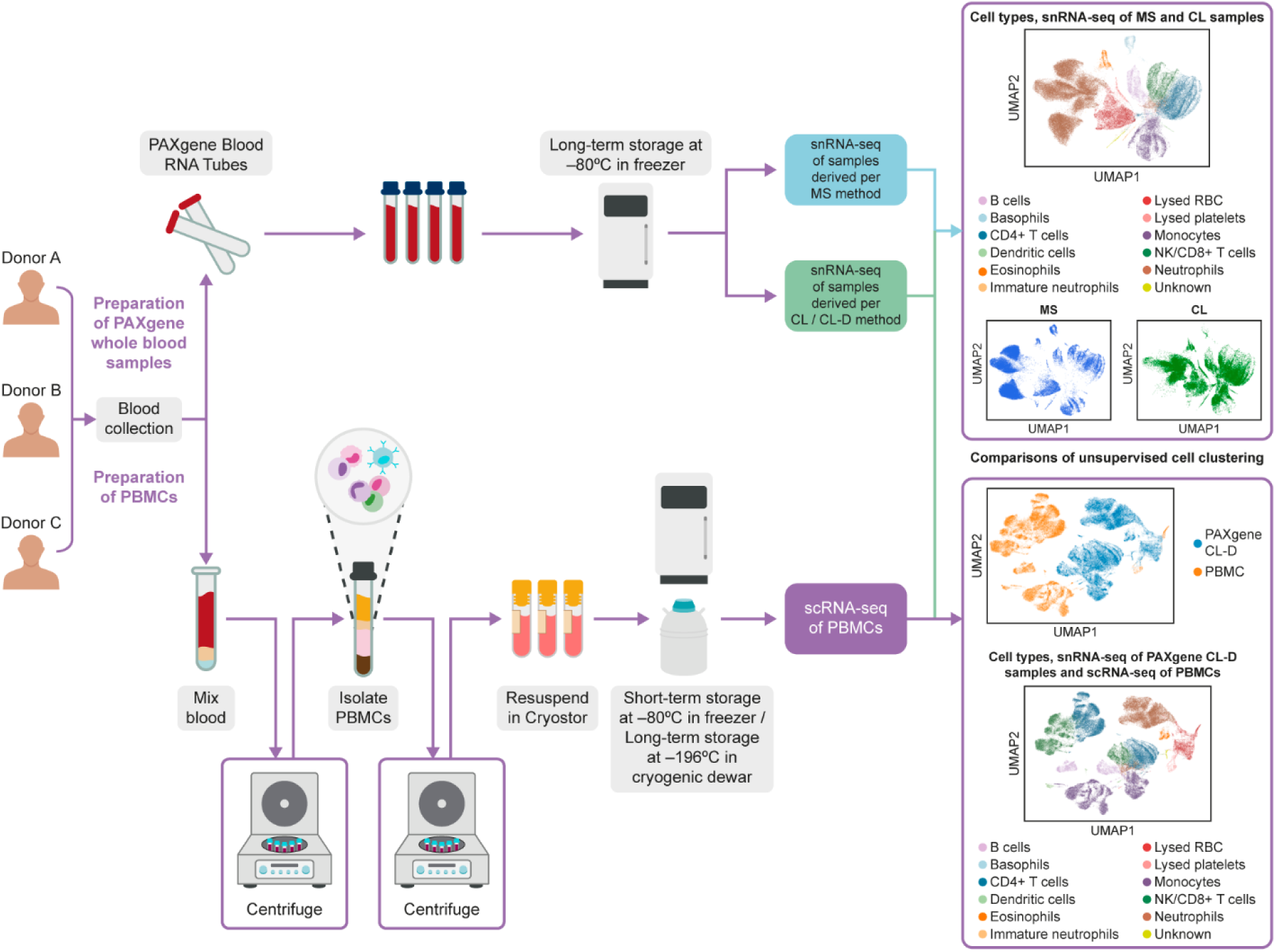
Whole blood collection, processing, nuclei isolation, and sequencing procedures. Fresh whole blood was processed using two procedures: PBMC extraction and cryopreservation followed by scRNA-seq, and PAXgene blood processing, freezing, and nuclei isolation (by mechanical separation or cell lysis) followed by snRNA-seq. CL(-D), cell lysis (with globin depletion); MS, mechanical separation; PBMC, peripheral blood mononuclear cell; scRNA-seq, single-cell RNA sequencing; snRNA-seq, single-nucleus RNA sequencing.

### Nuclei isolation methods for PAXgene whole blood samples

We developed two nuclei isolation methods: mechanical separation (MS) and cell lysis (CL). MS employed size exclusion and filter-adsorption to separate WBCs (10–20 µm diameter) from RBCs and debris (≤8 µm). The CL method involved pelleting whole blood, including RBCs, followed by cell lysis (see Methods, Figure S1B).

We assessed each method based on nuclei yield, sample composition, and the quality of the RNA captured (Figure 2A–C). Three main differences were observed. In terms of nuclei yield, MS consistently yielded fewer nuclei compared to CL. Across the three healthy donors, nuclei yield was up to 100-fold higher with CL (Figure 3A), indicating substantial nuclei loss during the MS procedure. With regards to sample composition, brightfield confocal microscopy images revealed higher amounts of debris following CL relative to MS (Figure S1C). Nuclear staining of samples from Donor A indicated that control (thawed PAXgene whole blood, Figure S1A) and CL-derived samples contained roughly equal proportions of single-lobed (lymphocytes) and multi-lobed (granulocytes) nuclei, whereas MS-derived samples had a higher proportion of multi-lobed nuclei (Figure 3B, Figure S1D). snRNA-seq analysis (Figure 3C) confirmed that the MS method resulted in samples that had significantly higher neutrophil proportions compared to CL-derived samples (ScanPro t-test, p = 0.026, Figure 3D). RNA library quality also differed between the methods. While complementary DNA (cDNA) yields for both methods were within the expected range of 250 bp to 2.5 kbp, traces for CL-derived samples (Figure S1E) showed a sharp peak attributed to abundant hemoglobin (globin) genes (*HBA1*, *HBA2*, *HBB*, *HBD*), likely from lysed RBCs (Figure 4A).

**Figure 2.**
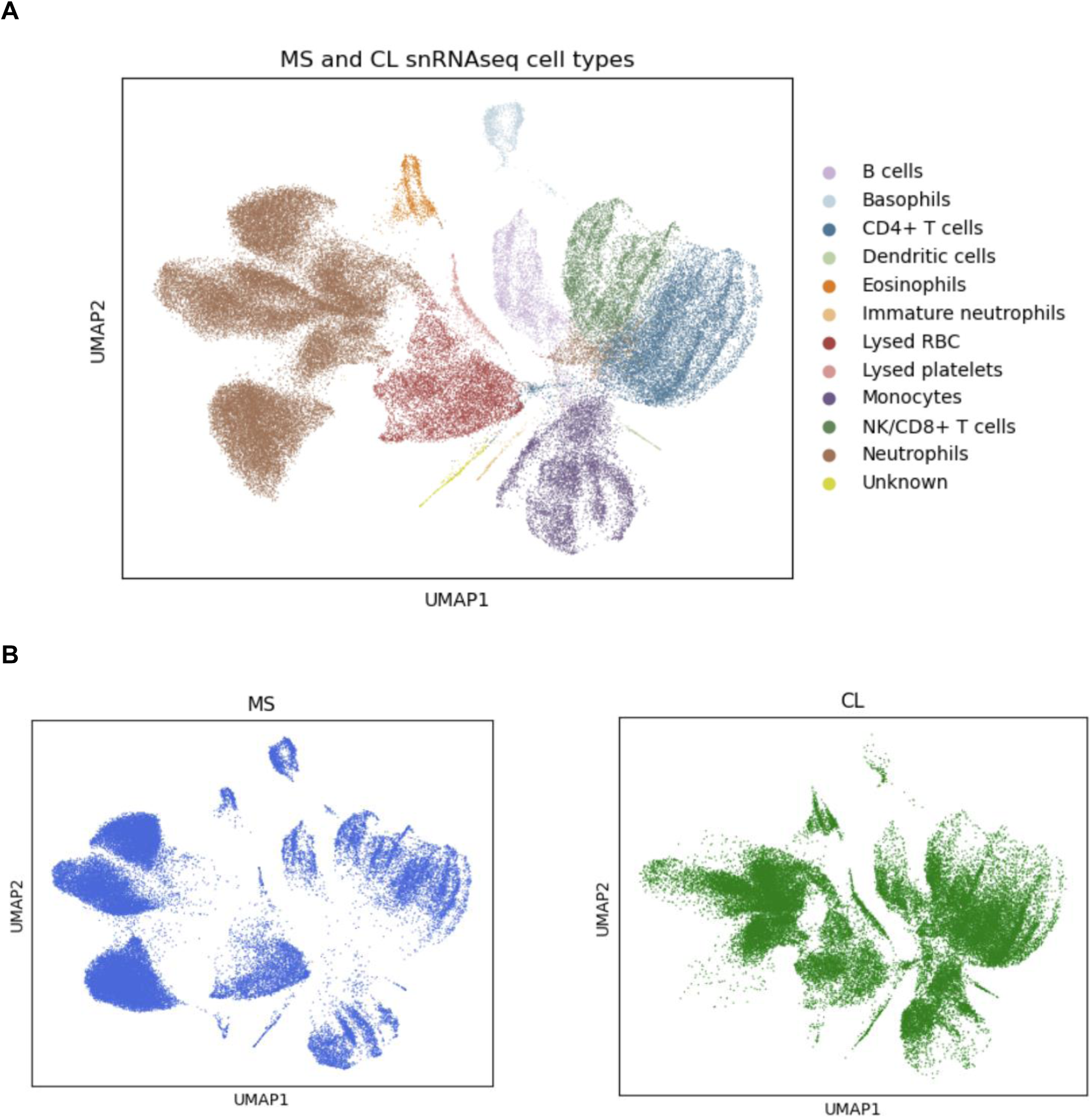
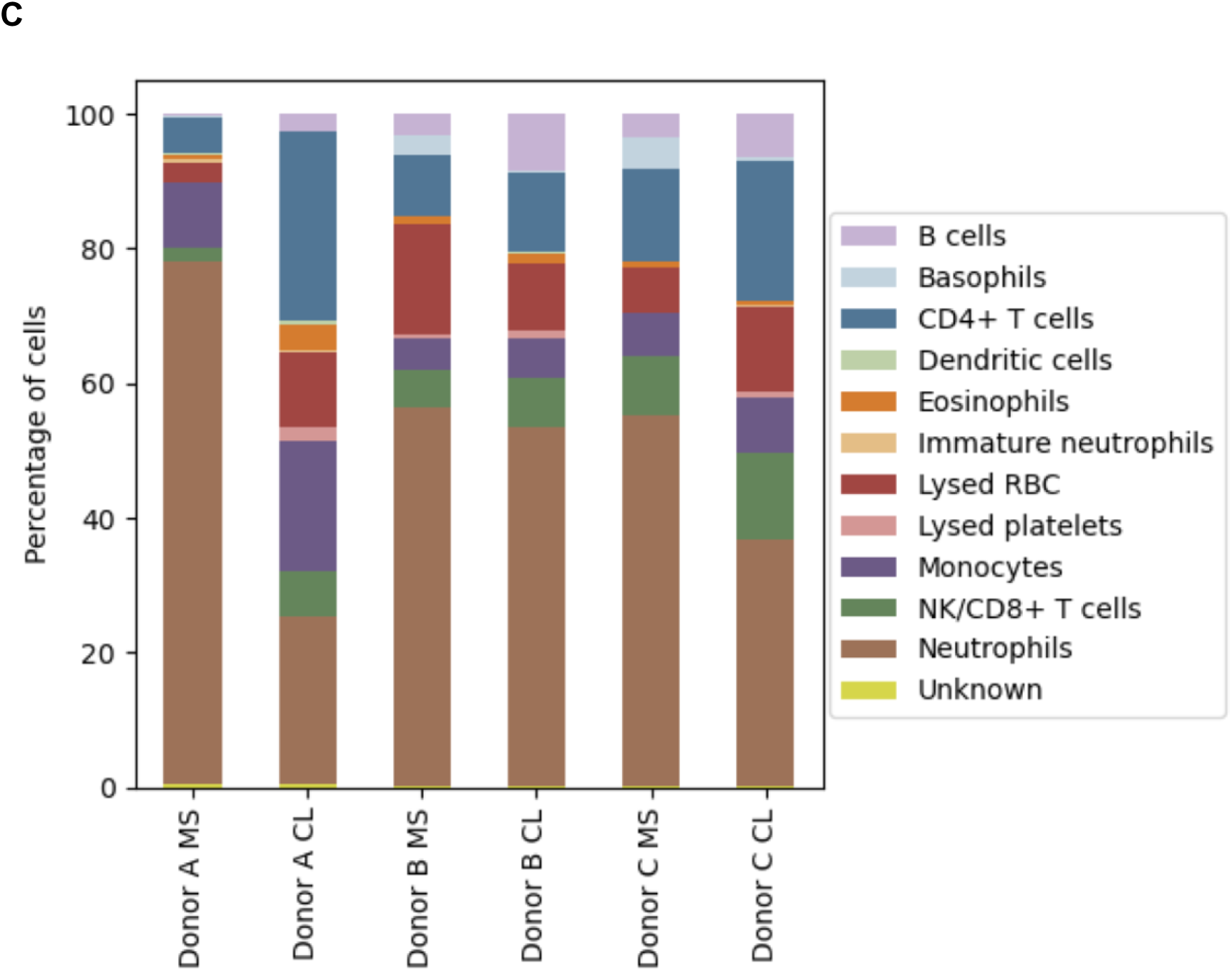
snRNA-seq analysis of nuclei isolated using the mechanical separation (MS) and cell lysis (CL) methods. (A) UMAP by cell type illustrates the detection of numerous cell types, including granulocytes (basophils, eosinophils, and neutrophils). (B) UMAP showing cell clusters for samples derived using the MS (blue) and CL (green) methods, based on analysis of duplicate samples per method from three donors (12 samples in total). (C) Proportions of cell types in samples derived using the MS and CL methods (bars show average of two duplicate samples per donor per method, 12 samples in total). CL, cell lysis; MS, mechanical separation; NK, natural killer; RBC, red blood cell; snRNA-seq, single-nucleus RNA sequencing; UMAP, uniform manifold approximation and projection.

**Figure 3.**
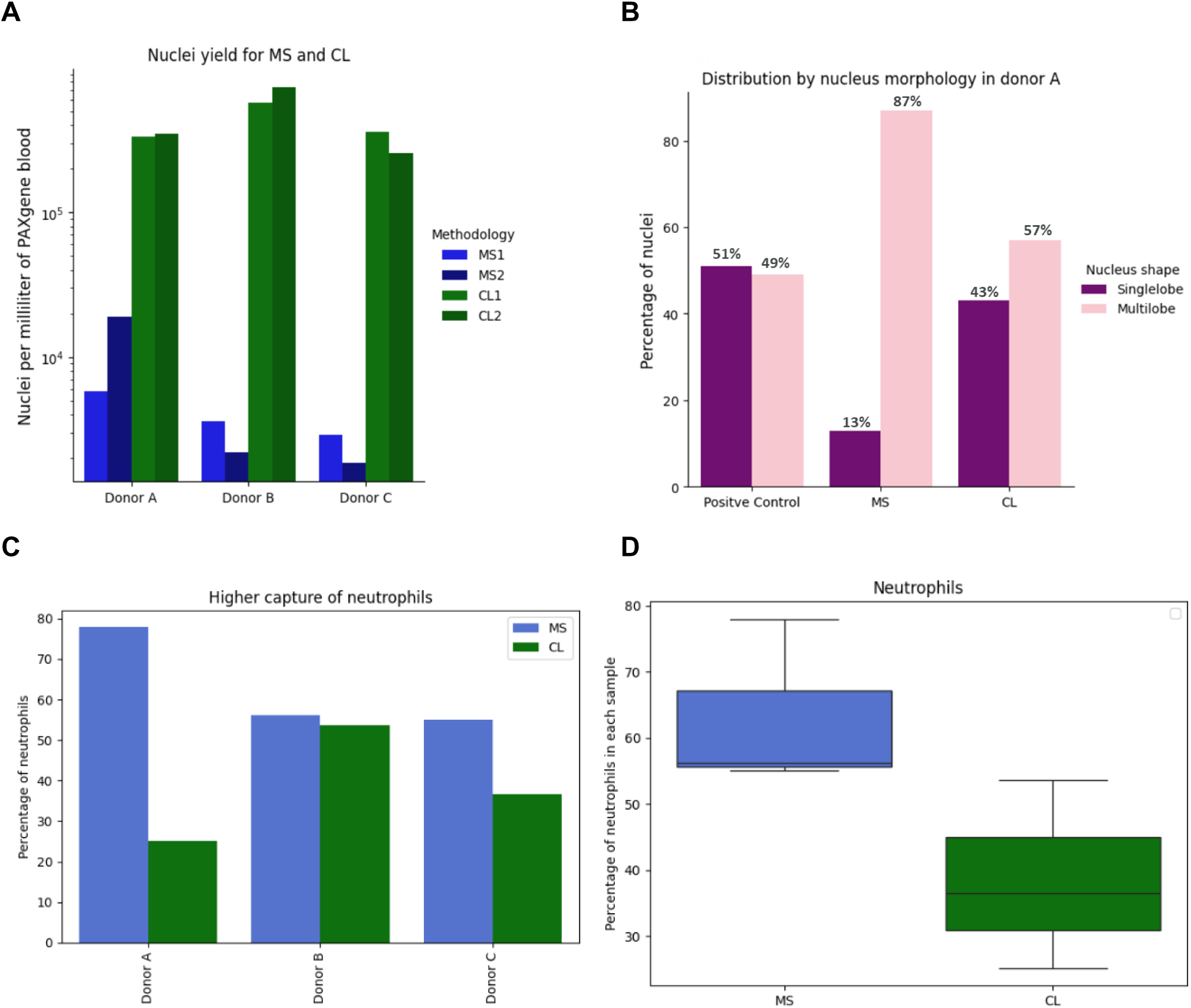
Nuclei yields and sample composition. (A) Comparison of nuclei yields per mL of PAXgene whole blood sample using mechanical separation (MS) and cell lysis (CL) nuclei isolation methods. The mean yields across the 6 samples (two duplicate samples per each of 3 donors) per method were 5,887 nuclei for MS and 434,746 nuclei for CL. (B) Composition in terms of nuclei morphology (multi-lobed versus single-lobed, by manual counting) of control (thawed PAXgene whole blood), MS-derived, and CL-derived samples. (C) MS-derived samples contained higher percentages of neutrophils compared to CL-derived samples (bars show average of two duplicate samples per donor per method, 12 samples in total). (D) Comparison of percentage of neutrophils in MS-derived versus CL-derived samples (t-test) demonstrates statistically significant enrichment of neutrophil populations with the MS method (6 samples, two duplicate samples per each of 3 donors). Horizontal line represents the median, box shows the interquartile range (IQR), and error bars represent 1.5 x IQR.

**Figure 4.**
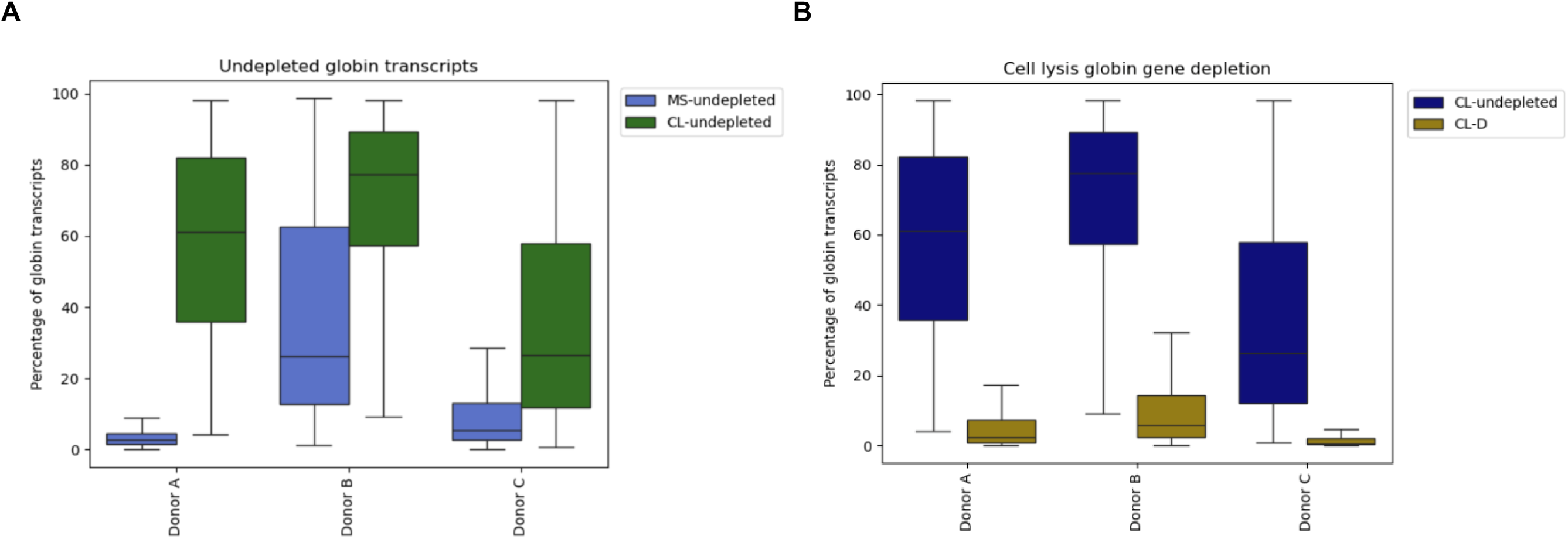
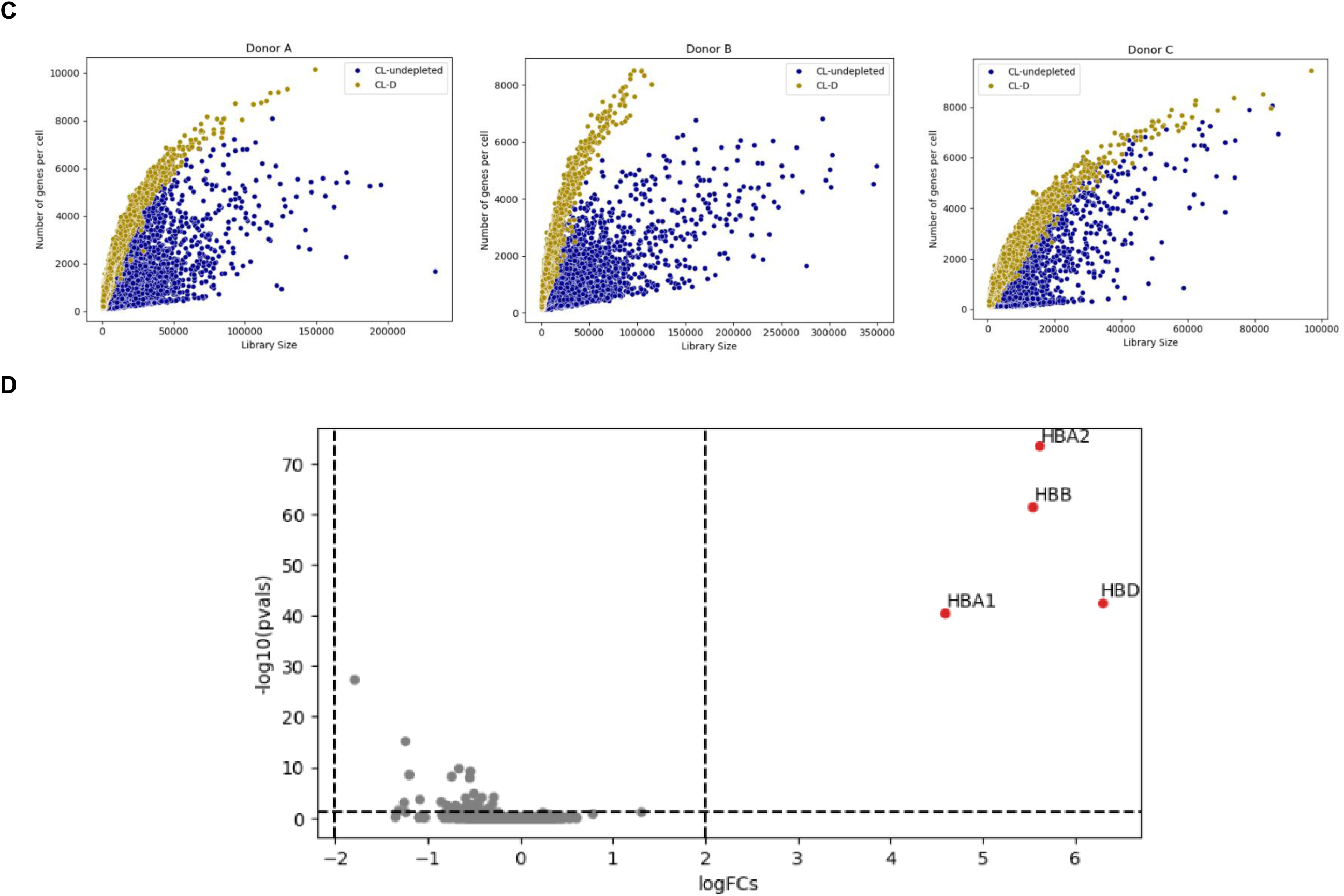
CRISPR-based globin transcript depletion to reduce RBC-derived ambient RNA. (A) Globin transcripts as percentages of total counts per cell for samples prepared by the mechanical separation (MS) and cell lysis (CL) methods, without depletion (MS-undepleted, CL-undepleted; 2 samples per box). Horizontal line represents the median, box shows the interquartile range (IQR), and error bars represent 1.5 x IQR. For comparison of pooled samples across donors prepared by MS and CL methods, t-test *p*-value < 0.0001. (B) Post-CL depletion of globin transcripts (CL-D method) per cell by CRISPR during final library construction resulted in substantial reductions in globin transcripts (2 samples per box). Horizontal line represents the median, box shows the IQR, and error bars represent 1.5 x IQR. For comparison of pooled CL-undepleted versus CL-D samples across donors, t-test *p*-value < 0.0001. (C) Individual nuclei data demonstrate an overall shift to higher numbers of genes per cell following globin depletion (dark mustard, CL-D method) relative to control (blue, CL method, no depletion; CL-undepleted). Library size is defined as total unique molecular identifier counts per cell. (D) Log2 fold change (FC) of individual gene expression between control (CL method, no depletion; CL-undepleted) and globin-depleted libraries (CL-D method) and associated –log10 *p*-values (duplicate samples from each of the 3 healthy donors, 6 samples in total).

### Ambient globin RNA depletion using CRISPR

The CL method was preferred to the MS method due to producing higher nuclei yields and more proportional WBC composition (Figure 2A–C); however, mean RNA contamination of 37.6–71.3% globin transcripts for the three donors (Figure 4A) needed to be reduced by CRISPR-based depletion^8,9^ of the four globin genes *HBA1*, *HBA2*, *HBB*, and *HBD* (Figure S2). CRISPR-treated samples showed significant globin depletion relative to control samples (t-test *p*-value < 0.0001, Figure 4B), and globin-depleted samples had more genes detected per cell than undepleted samples (Figure 4C), indicating higher sequencing depth and saturation. Analysis of pseudobulk data found that the only differentially expressed genes (with a log2 fold change of > 2 or < –2, adjusted *p*-value < 0.05) between globin-depleted and non-globin-depleted libraries were the target globin genes (Figure 4D). Globin transcript depletion was consistent across cell types (Figure S3).

### Whole blood snRNA-seq complements PBMC scRNA-seq

We compared snRNA-seq data from nuclei isolated from PAXgene whole blood samples using the CL plus globin-depletion (CL-D) method with scRNA-seq data from matched PBMC samples (Table 1). As anticipated, nuclei isolated using the CL-D method clustered separately from PBMC cells due to inherent differences in nuclear versus whole-cell transcripts (Figure 5A,B). However, we found similar proportions of mononuclear cell types in both groups (Figure 5C,D). Only the proportions of B cells (*p* = 0.04) and of dendritic cells (*p* = 0.04) were significantly different between PBMCs and CL-D-isolated nuclei (Figure 5E), and these cells constituted low percentages of the overall cell population (14% and 0.1% of all mononucleated cells, respectively; Figure 5C).

**Figure 5.**
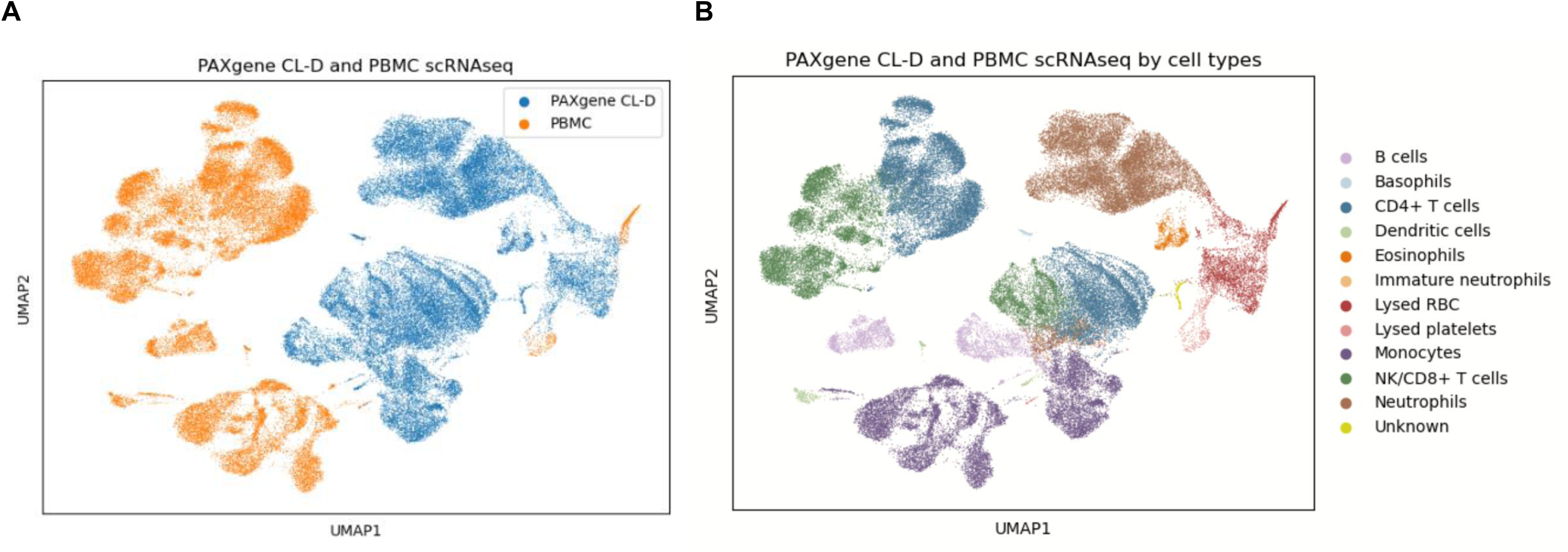
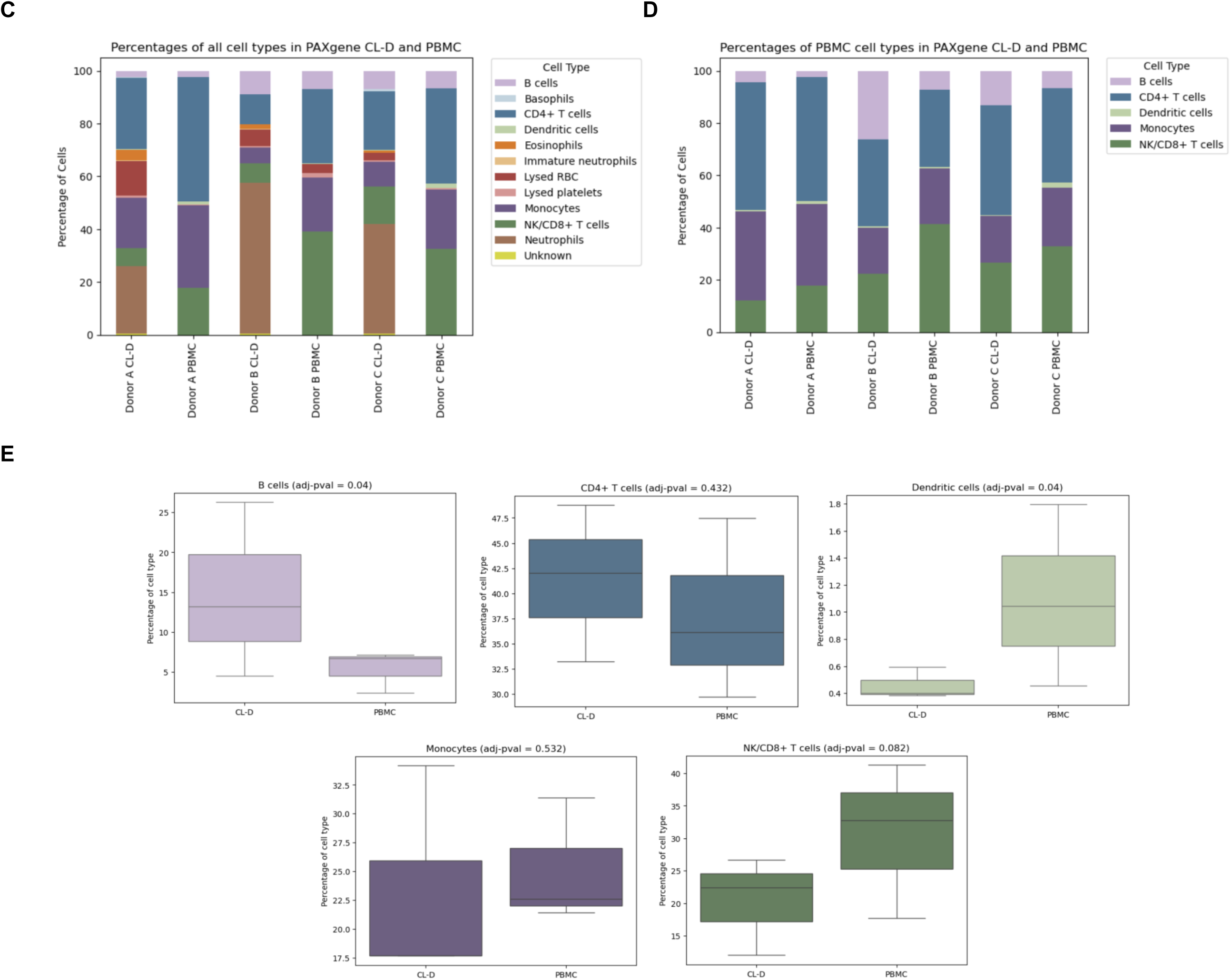
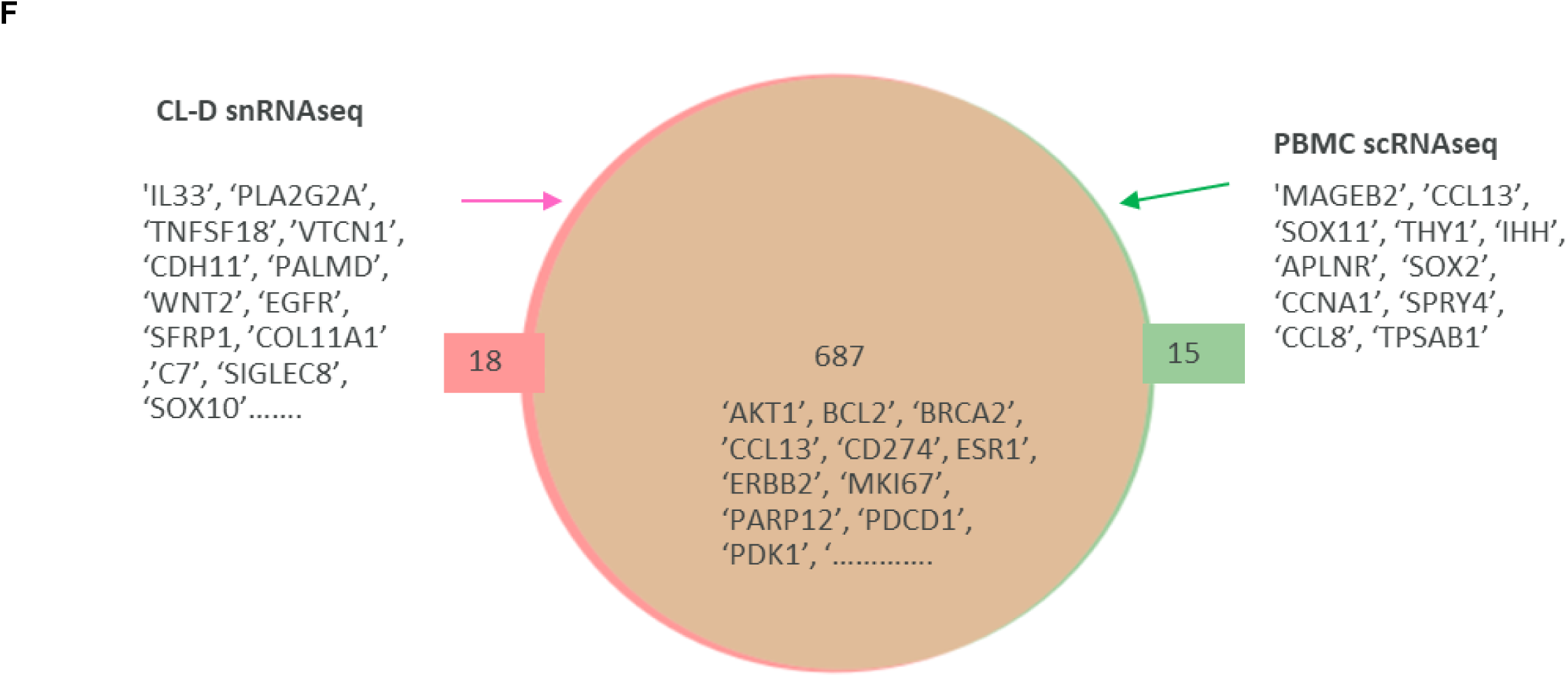
Comparison of unsupervised cell clustering of snRNA-seq data from PAXgene whole blood samples (CL-D nuclei isolation method) and scRNA-seq data from matched PBMC samples. (A) UMAP representation showing cell clusters derived from snRNA-seq data for nuclei isolated from PAXgene whole blood samples using the CL-D method (blue) and scRNA-seq data for matched PBMC samples (orange), using duplicate samples per donor per method (12 samples in total). (B) UMAP representation colored according to cell type. Although clusters were separate according to preparation method, clusters of similar cell types were identified by both methods, with the exception of granulocyte clusters (brown, orange, light orange, light blue) and lysed RBC (red) and lysed platelets (pink) and unknown (yellow) that were unique to snRNA-seq data for PAXgene whole blood samples. (C, D) 100% stacked bar plots of cell-type composition showing (C) all cell types and (D) cell types excluding granulocytes identified from PAXgene whole blood samples and matched PBMC samples (average of duplicate samples per donor per method; 12 samples in total). (E) Comparisons of individual cell type percentages between snRNA-seq data for PAXgene whole blood samples and scRNA-seq data for matched PBMC samples (*p*-values from Scanpro significance test; average of duplicate samples from 3 donors per method). Horizontal line represents the median, box shows the interquartile range (IQR), and error bars represent 1.5 x IQR. (F) Venn diagram of selected nCounter^®^ PanCancer IO 360™ Panel genes showing detection ability for snRNA-seq of PAXgene whole blood samples and scRNA-seq of matched PBMC samples (duplicate samples from 3 donors per method; 6 samples in total). Gene count filter was 5 for detection. CL-D, cell lysis with globin depletion; MS, mechanical separation; NK, natural killer; PBMC, peripheral blood mononuclear cell; RBC, red blood cell; scRNA-seq, single-cell RNA sequencing;snRNA-seq, single-nucleus RNA sequencing; UMAP, uniform manifold approximation and projection.

To confirm that clinically relevant genes were captured by snRNA-seq of nuclei isolated from PAXgene whole blood samples using the CL-D method, we compared gene detection by snRNA-seq in CL-D-derived samples and scRNA-seq in PBMC samples using a reference set of immune-related genes (Nanostring IO360 panel, (https://nanostring.com/products/ncounter-assays-panels/oncology/pancancer-io-360/).^10^ Out of 751 common genes, 687 genes were detected by both methods, 18 were detected only on snRNA-seq of CL-D derived PAXgene whole blood samples, 15 were detected only on scRNA-seq of PBMC samples, and 31 were not detected by either method (Figure 5F). Gene detection sensitivity was not compromised even though CL-D-derived samples contained nuclei rather than intact cells.

## DISCUSSION

PAXgene RNA Blood Tubes are commonly used for the rapid collection and preservation of whole blood samples, and thousands of these samples are typically collected in large biobanking and biopharmaceutical research studies. Previously, the nature of frozen, preserved whole blood samples precluded single-cell transcriptomic analysis. This manuscript describes the novel CL-D method for isolating intact nuclei from PAXgene whole blood samples for snRNA-seq analysis.

Initial comparisons showed that the CL nuclei isolation method resulted in a substantially higher nuclei yield than the MS method. Additionally, the proportions of cell types in CL-derived samples more closely matched the proportions seen in control PAXgene whole blood samples that had not undergone nuclei isolation. Further differences between the methods are summarized in Table 2. The main drawback to the CL method was that ambient RNA was left over from the lysis process; this was resolved using CRISPR-based removal of ambient RNA globin gene transcripts. snRNA-seq data for samples derived using the CL-D method, incorporating CL and CRISPR-depletion (D), compared favorably to scRNA-seq data from matched PBMC samples, with similar cell type proportions, gene expression, and gene sensitivity. CL-D-derived samples had the additional benefit of including granulocytes, which have an important role in the innate immune response.^11^ Neutrophils have been used as predictive markers in disease progression,^12–14^ patient survival in breast and lung adenocarcinomas,^15^ sepsis,^16^ and metabolic syndrome.^17^ Neutrophil subtypes can have pro-tumor or anti-tumor characteristics, and, under certain circumstances, neutrophils may change from one subtype to another.^18,19^

**Table 2.** Head-to-head comparison of the two nuclei isolation methods for processing PAXgene whole blood samples for snRNA-seq, highlighting key differences and unique features of each method.

|  | Mechanical separation (MS) | Cell lysis (CL) |
| --- | --- | --- |
| **Sample input** | 4 mL to 9 mL | 4 mL is sufficient |
| **Nuclei yield** | Unreliable and low | Reliable and high |
| **Nuclei morphology bias** | Enrichment of granulocytes | Not apparent |
| **Composition reproducibility** | Mononuclear: mid-high<br>Granulocytes: mid | Mononuclear: high<br>Granulocytes: mid |
| **Sample preparation** | Longer (tricky) | Straightforward |
| **Single session throughput** | 2 samples | 4 samples |
| **Data quality at baseline** | High | Mid |
| **Data quality post globin depletion** | High | High |
| **Bulk RNA option** | Unlikely | Yes |
| **Cost** | Similar | Similar |

snRNA-seq, single-nucleus RNA sequencing.

Our methodology focused on isolating nuclei from preserved whole blood in an effort to enable RNA-seq with single-cell resolution for these samples, for which previously only bulk analyses were possible. We tested whole blood samples preserved in PAXgene tubes specifically due to the widely established use of these tubes and the ease of specimen collection. PAXgene tubes are manufactured with all reagents included and keep RNA stable at ambient temperature for up to 3 days. Other commercial methods^20,21^ for preserving whole blood involve proprietary equipment,^21^ techniques^22,23^, and/or additional steps such as flow cytometry.^3^

Our CL-D method for isolating nuclei could also be tested for use with other blood collection tubes such as PAXgene Blood DNA Tubes, Tempus blood RNA tubes, *Quick*-DNA/RNA tubes (Zymo Research), and Blood STASIS tubes (MAGBIO Genomics). Samples processed using our CL-D method can likely be sequenced on single-cell platforms other than those from 10x Genomics that were used in these analyses, such as platforms from Parse Biosciences, Scale Biosciences, and Fluent BioSciences. Our novel method, leveraging easy blood collection at medical sites, has the potential to unlock many underutilized frozen whole blood samples and provide insights into the biology of all immune cell types.

### Limitations of the study

While we demonstrate that CL-D snRNA-seq is applicable to PAXgene Blood RNA Tubes using various aliquots from three healthy donors, profiling samples from additional donors will confirm the robustness of the presented method. Furthermore, ground truth of donor blood composition via complete cell count (CBC) or fluorescence-activated cell sorting (FACS) counting was not available, and future studies with such validation experiments will confirm the accuracy of detected cell types by our isolation and sequencing methodologies. It should be noted that fresh blood is required for both CBC and FACS counting, and so these methods are incompatible with PAXgene preserved blood samples. Finally, while our methodology tested PAXgene preserved blood, the CL-D method may be tested and adapted for alternative preserved blood collection methods, including the ones mentioned in the Discussion section.

Nevertheless, we have shown that our method allows researchers to explore single-cell transcriptomics of circulating blood using a simplified procedure.

## RESOURCE AVAILABILITY

### Lead contact

Further information and requests for resources and reagents should be directed to and will be fulfilled by the lead contact, Ojasvi Chaudhary.

### Data and code availability

RNA sequencing data underlying the findings of this manuscript may be obtained in accordance with AstraZeneca’s data sharing policy, per the information provided in the ‘Lead contact’ section. This paper does not report original code. Any additional information is available from the lead contact upon request.

## ACKNOWLEDGMENTS

The authors thank the following colleagues from AstraZeneca: Elza De Bruin and Dhivya Sudhan for input and feedback, and Minoo Rafati for early project work, and Megan Callahan and William Kelly for support in preparation of the manuscript. Medical editing support for the development of this manuscript, under the direction of the authors, was provided by Steve Hill, PhD, of Ashfield MedComms (Macclesfield, UK), an Inizio company, and was funded by AstraZeneca.

## AUTHOR CONTRIBUTIONS

**Conceptualization:** O.C., G.D., and A.R.

**Methodology:** O.C., G.D., P.G., R.A., and M.R.

**Validation:** X.C., E.G., and R.H.

**Formal analysis:** M.Steinberg and G.D.

**Data Curation:** M.Steinberg, V.S., and C.R.

**Writing - Original Draft:** O.C., M.R., and A.R.

**Writing - Review & Editing:** All authors.

**Visualization:** G.D. and A.R.

**Supervision:** B.D., A.R., and J.B.

**Project administration:** P.G.

**Funding acquisition:** B.D. and M.Scaltriti

## FUNDING

These analyses were funded by AstraZeneca.

## DECLARATION OF INTERESTS

Dr. Maurizio Scaltriti is employed by AstraZeneca, owns AstraZeneca stock and/or options, and is a co-founder of Medendi.

Rogelio Aguilar was employed by AstraZeneca at the time of this study.

The remaining authors are employed by AstraZeneca and own AstraZeneca stock and/or options; they declare no other competing interests.

