## Supplemental Information for "Comprehensive cellular analysis with single-nucleus RNA-seq of archived PAXgene whole blood samples"

### KEY RESOURCES TABLE

| REAGENT OR RESOURCE | SOURCE | IDENTIFIER |
|---|---|---|
| **Biological material** | | |
| Healthy donor blood | Research Blood Components (Watertown, MA) | https://www.researchbloodcomponents.com/ |
| **Commercial assays and systems** | | |
| BD vacutainer CPT mononuclear cell preparation tubes | BD Biosciences | Cat. No. 362753 |
| CoolCell LX freezing containers | Corning | Cat 432138 |
| BD vacutainer K2 EDTA tubes | Fisher Scientific | Cat. 02-657-32 |
| PAXgene® blood RNA tubes | Qiagen | Cat. 762165 |
| Acrodisc syringe filters | Pall Corporation | Cat. AP-4952 |
| 100 µm cell strainer | Corning | Cat. 431752 |
| 40 µm cell strainer | Corning | Cat. 431750 |
| Cellometer Auto 2000 | Nexcelom | CMT-A2K |
| 3’ Kit v3.1(Chip G) | 10x Genomics | CG000315 Rev E |
| NovaSeq 6000 system | Illumina | 3376672 |
| DepleteX™ | Jumpcode Genomics | Cat. KIT1024 |
| IO360 panel | Nanostring | https://nanostring.com/products/ncounter-assays-panels/oncology/pancancer-io-360/ |
| **Chemicals** | | |
| 10% Fetal Bovine Serum (FBS) | Thermofisher scientific | Cat. A3160401 |
| Dulbecco’s Phosphate Buffered Saline (DPBS) | Thermofisher scientific | Cat. 14190144 |
| CryoStor | Stemcell Technologies | Cat. 100-1061 |
| 10 % MACS BSA | Miltenyi Biotec | Cat. 130-091-376 |
| RNase inhibitor murine | New England BioLabs | Cat. M0314L |
| UltraPure distilled water | Thermofisher scientific | Cat. 10977015 |
| Trizma hydrochloride solution 1M | Millipore sigma | Cat. T2194 |
| Sodium chloride solution 5M | Millipore sigma | Cat. 59222C |
| Magnesium chloride solution 1M | Millipore sigma | Cat. M1028 |

| REAGENT OR RESOURCE | SOURCE | IDENTIFIER |
|---|---|---|
| 10% Tween-20 | Bio-Rad | Cat. 1662404 |
| Nonidet P40 substitute | Millipore sigma | Cat. 74385 |
| IGEPAL CA-630 | Millipore sigma | Cat. I8896 |
| 5% Digitonin | Thermofisher scientific | Cat. BN2006 |
| Dithiothreitol (DTT) 1M | Millipore sigma | Cat. 646563 |
| Acridine orange and propidium iodide | Revvity | CS2-0106-25ML |
| SPRIselect | Beckman Coulter | Cat. B23318 |
| Software/Data resources | | |
| 10x Cell Ranger (6.1.1) | 10x Genomics | https://support.10xgenomics.com/single-cell-gene-expression/software/pipelines/ |
| scanpy (v1.10.1) | Wolf et al, 2018[23] | https://github.com/scverse/scanpy |
| scrublet (v0.2.3) | Wolock et al, 2019[24] | https://github.com/swolock/scrublet |
| Leiden graph-clustering method | | |
| Scanpro (v0.3.2) | Alayoubi et al, 2024[25] | https://github.com/loosolab/scanpro |
| decoupleR (v1.5.0) | Badia-i-Mompel et al, 2022[26] | https://github.com/saezlab/decoupler-py |
| pyDESeq2 (v0.4.10) | Muzellec et al, 2023[27] | https://github.com/owkin/PyDESeq2 |

### METHOD DETAILS

#### Isolation of peripheral blood mononuclear cells (PBMCs)

Whole blood samples from three healthy individual donors (Donors A, B, and C) were acquired from Research Blood Components (Watertown, MA), and aliquots were collected in BD Vacutainer CPT mononuclear cell preparation tubes (BD Biosciences). Cells were washed in 10% Fetal Bovine Serum (FBS) in Dulbecco's Phosphate Buffered Saline (DPBS) without calcium and magnesium. CPT tubes containing whole blood and sodium citrate were gently mixed at room temperature (RT) and then underwent centrifugation at 1600 x g for 20 min without braking. The white PBMC layer beneath the yellow plasma layer was collected using a 5 mL serological pipette and transferred to a 15 mL conical tube. PBMCs were washed twice with 10% FBS at 300 x g for 10 min (first wash) and 5 min (second wash). A small aliquot was used for cell counting after the first wash; then, after the second wash, the supernatant was removed and the pellet resuspended in CryoStor (Stemcell Technologies) at a concentration of ≈ 2 × $10^6$/mL. Subsequently, cells were frozen (1 mL per cryovial) at –1ºC per minute in Corning CoolCell LX freezing containers (Figure 1).

### Preparation of whole blood samples in PAXgene RNA Blood Tubes

Aliquots of the same whole blood samples used for PBMC isolation were collected in BD Vacutainer K2 EDTA tubes (Fisher Scientific) and gently mixed; 2.5 mL of blood was added to individual PAXgene tubes (Qiagen) containing 6.9 mL of preservative. Tubes were stored at –80ºC (Figure 1), at which temperature the samples remain stable for up to 11 years.[5] Each whole blood sample was processed in duplicate to confirm repeatability of results.

### Isolation of nuclei from PAXgene whole blood samples

Nuclei suspensions were prepared from whole blood samples stored in PAXgene RNA Blood Tubes via two methods (Figure S1B) – mechanical separation (MS) and cell lysis (CL). Frozen PAXgene RNA Blood Tubes that had been stored at –80ºC were slowly thawed overnight at 4ºC then equilibrated at RT for 2 hr. Based on our own experience regarding the maximum number of samples processed by one person at a time, is recommended that an individual user does not process more than 4 samples simultaneously. Subsequently, the two nucleus isolation procedures below were implemented.

#### *MS methodology*

A wash buffer of PBS + 0.04% bovine serum albumin (BSA) + 0.04% RNase inhibitor, murine (NEB) was prepared. Three Acrodisc syringe filters (Pall Corporation) were mounted on 5 mL syringes, with the Pall logo facing the plunger, and placed on top of 50 mL Falcon tubes. The three filters were each primed with 5 mL wash buffer, and ≈3 mL of mixed PAXgene whole blood at RT was added to each filter and allowed to flow through by gravity; the contents of a single PAXgene Blood RNA Tube were equally distributed across the three filters. After 3–5 min, the filters were washed with 5 mL wash buffer. Each Acrodisc was then unscrewed, flipped, and attached to a new 5 mL syringe; 5 mL wash buffer was plunged through the Acrodisc to dislodge and collect nuclei from the filter (Leukosorb membrane). Nuclei from each filter were pooled into one 15 mL tube, washed with wash buffer, and centrifuged at 500 x g for 5 min at 4ºC. The pellet was filtered through a 100 µm cell strainer and then a 40 µm cell strainer (Corning) and centrifuged at 500 x g for 5 min at 4ºC. The nuclei were resuspended in ≤500 µL of wash buffer and the suspension kept on ice.

#### *CL methodology*

A lysis buffer was prepared per protocol CG000365 Rev C (10x Genomics) comprising UltraPure distilled water, 10 mM Tris-HCl, pH 7.4, 10 mM NaCl, 3 mM $MgCl_2$, 0.1% Tween-20, 0.1% Nonidet P40 substitute or IGEPAL CA-630, 0.01% digitonin, 1% BSA, 1 mM DTT, and RNase inhibitor, murine. Similarly, a lysis wash buffer was prepared per protocol CG000365 Rev C comprising UltraPure distilled water, 10 mM Tris-HCl, pH 7.4, 10 mM NaCl, 3 mM $MgCl_2$, 0.1% Tween-20, 1% BSA, 1 mM DTT, and RNase inhibitor, murine, together with wash buffer and water plus RNase inhibitor (0.04% RNase inhibitor, murine [NEB] plus ultra-distilled water).

PAXgene whole blood samples at RT were mixed by inverting and resuspending with a 10 mL serological pipette. After mixing, 4 mL of blood was transferred to a 15 mL Falcon tube and centrifuged at 3000 x g for 10 min at RT. The supernatant was removed, and the large red pellet was resuspended in 4 mL water plus RNase inhibitor and centrifuged again at 3000 x g for 10 min at RT. The supernatant was removed, and the dark pellet was lysed in 350 µL lysis buffer for 5 min on ice. Immediately afterward, 3.5 mL of lysis wash buffer was added to stop lysis. The sample was filtered through a 100 µm cell strainer and then a 40 µm cell strainer. The filtered nuclei were washed in wash buffer and centrifuged. The pelleted nuclei were resuspended in ≥500 µL of wash buffer and kept on ice.

#### Single-nucleus RNA sequencing (snRNA-seq)

Portions of nuclei suspensions derived using the MS and CL methods were stained with acridine orange and propidium iodide (AO/PI) and counted on a Cellometer Auto 2000 (Nexcelom). Sufficient nuclei were loaded onto Next GEM Chip G chip (10x Genomics) per the manufacturer protocol CG000315 Rev E, targeting 5000 nuclei for single-nucleus libraries. Libraries were sequenced on the Illumina NovaSeq 6000 system.

#### CRISPR-guided RNA depletion of CL-derived nuclei samples

Single-cell barcoding, reverse transcription, generation of double-stranded DNA from single-stranded cDNA, and gene expression library construction were performed per protocol CG000315 Rev E (10x Genomics). The CRISPRclean RNA Depletion (Globin) kit, now called DepleteX (Jumpcode Genomics), was used to deplete globin genes (*HBA1*, *HBA2*, *HBB*, and *HBD*). For optimal results, 20–200 ng of cDNA input and 12 index PCR cycles were used. The ribonucleoprotein (RNP) complex was prepared during ligation reaction and incubated at RT for 10 min. For post-ligation cleanup, Cas9 buffer and RNP complex was added to the sample, which was then incubated for 60 min at 37ºC. After incubation the reaction was immediately transferred to ice for 10 min, followed by 0.6X SPRI selection with a final elution volume of 30 µL. Index PCR added i5 and i7 barcodes for sample identification post-sequencing, which was conducted on an Illumina NovaSeq 6000. Sequencing parameters were consistent with protocol CG000315 Rev E (10x Genomics). Depleted libraries had longer average sizes than control undepleted libraries due to the removal of short repetitive globin transcripts (Figure S1D).

### QUANTIFICATION AND STATISTICAL ANALYSIS

#### Single-cell and single-nucleus data analysis

The cellranger mkfastq pipeline (version 6.1.1; 10x Genomics) was used to demultiplex the base call files (BCLs) to FASTQ files, and then the cellranger count pipeline was used for downstream processing steps, including alignment to reference genome GRCh38, counting of unique molecular identifiers (UMIs), and cell calling. Gene expression matrices were analyzed using scanpy[23] (v1.10.1) best practices.[28] Quality filters removed cells with fewer than 100 molecules, genes found in fewer than three cells, and cells with >20% mitochondria. Doublets were identified using scrublet[24] (v0.2.3), and cells with scores >0.72 were removed.

Remaining cells were normalized to counts per million and log-transformed. We identified and scaled 2421 highly variable genes based on mean expression (0.0125–3) and dispersion >0.5. These genes were used to calculate principal components, which were used to create a Uniform Manifold Approximation and Projection (UMAP) and embed the K-Nearest Neighbor (KNN) graph. The KNN graph was clustered with the Leiden graph-clustering method.

Cell types were annotated using known marker genes (Table S1). When comparing subsets of the full dataset (e.g., CL-derived samples only), the samples of interest were reanalyzed to find new principal components, and new clusters and UMAPs were generated for the subset. Proportions of cell types were compared between methods (e.g. MS vs. CL) using Scanpro[25] (v0.3.2). Pseudobulk profiles were created using decoupleR[26] (v1.5.0), and differential expression analysis between methods was conducted using pyDESeq2[27] (v0.4.10).

**Table S1. Marker genes used to identify cell type clusters in PAXgene snRNA-seq and PBMC scRNA-seq data**

| Cell type | Marker genes |
|---|---|
| **B cells** | CD79A+, MS4A1+ |
| **Basophils** | CCR3+, ENPP3+ |
| **CD4+ T cells** | CD3E+, CD4+ |
| **Dendritic cells** | IL3RA+, CLEC4C+, NRP1+ |
| **Eosinophils** | CCR3+, ADGRE1+ |
| **Immature neutrophils** | CEACAM8+, NKG7+, DEFA3+ |
| **Monocytes** | CD14+, FCGR3A+ |
| **NK / CD8+ T cells** | CD3E+, CD8+, NKG7+ |
| **Neutrophils** | FCGR2A+, ITGAX+ |
| **Lysed Platelets** | PPBP+, CLU+, PF4+, F13A1+ |
| **Lyesed Red blood cells** | HBA1+ |

**Figure S1. Processing and imaging of PAXgene whole blood samples**

(A) Confocal microscopy images of a blood pellet from a PAXgene whole blood sample resuspended in PBS + 0.04% BSA. Scale bar is 50 µm. (B) Schematic illustration of mechanical separation (MS) and cell lysis (CL) nuclei isolation methods. (C) Confocal microscopy images of nuclei suspensions following nuclei isolation using the MS and CL methods. Scale bar is 50 µm. (D) Confocal microscopy images highlighting single-lobed (T cells, B cells, dendritic cells) and multi-lobed (granulocytes) cell types in an MS-derived sample. (E) Complementary DNA (cDNA) amplified product and final library traces for both the MS and CL nuclei isolation methods (representative sample from Donor A).

**A**

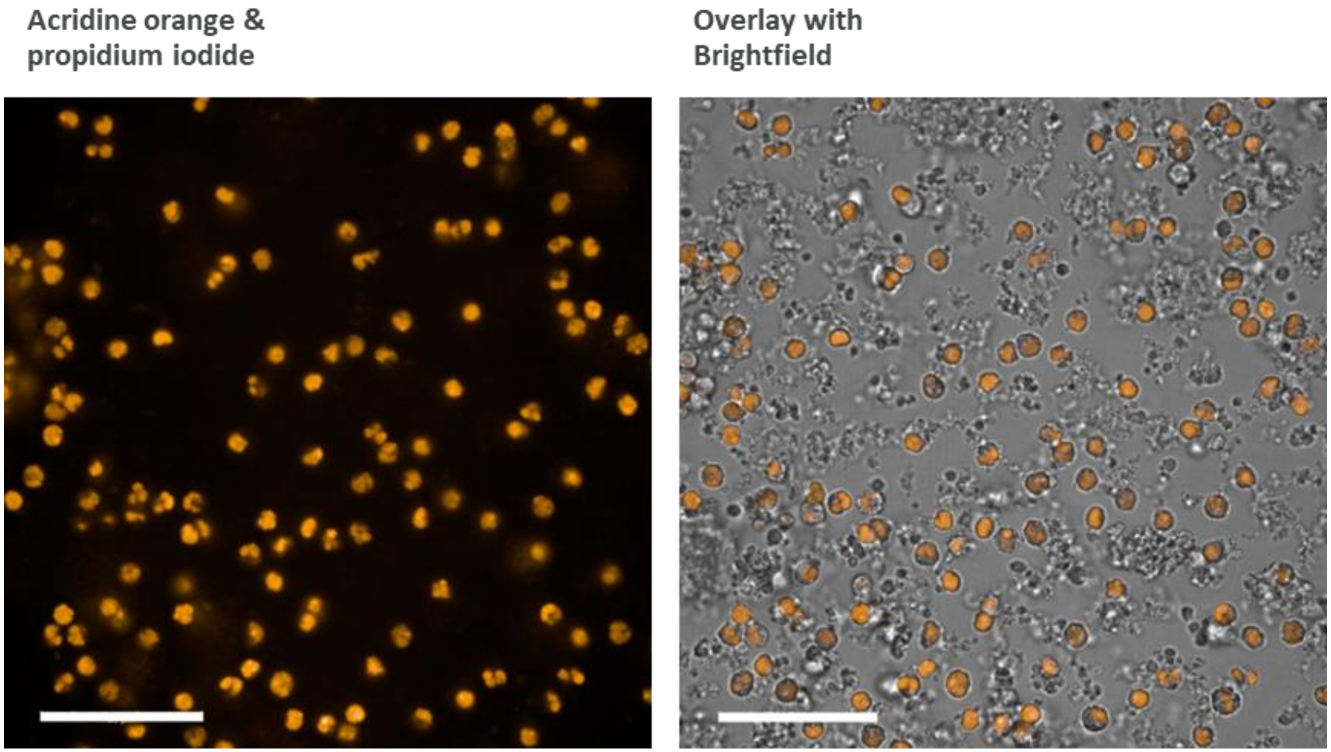

B

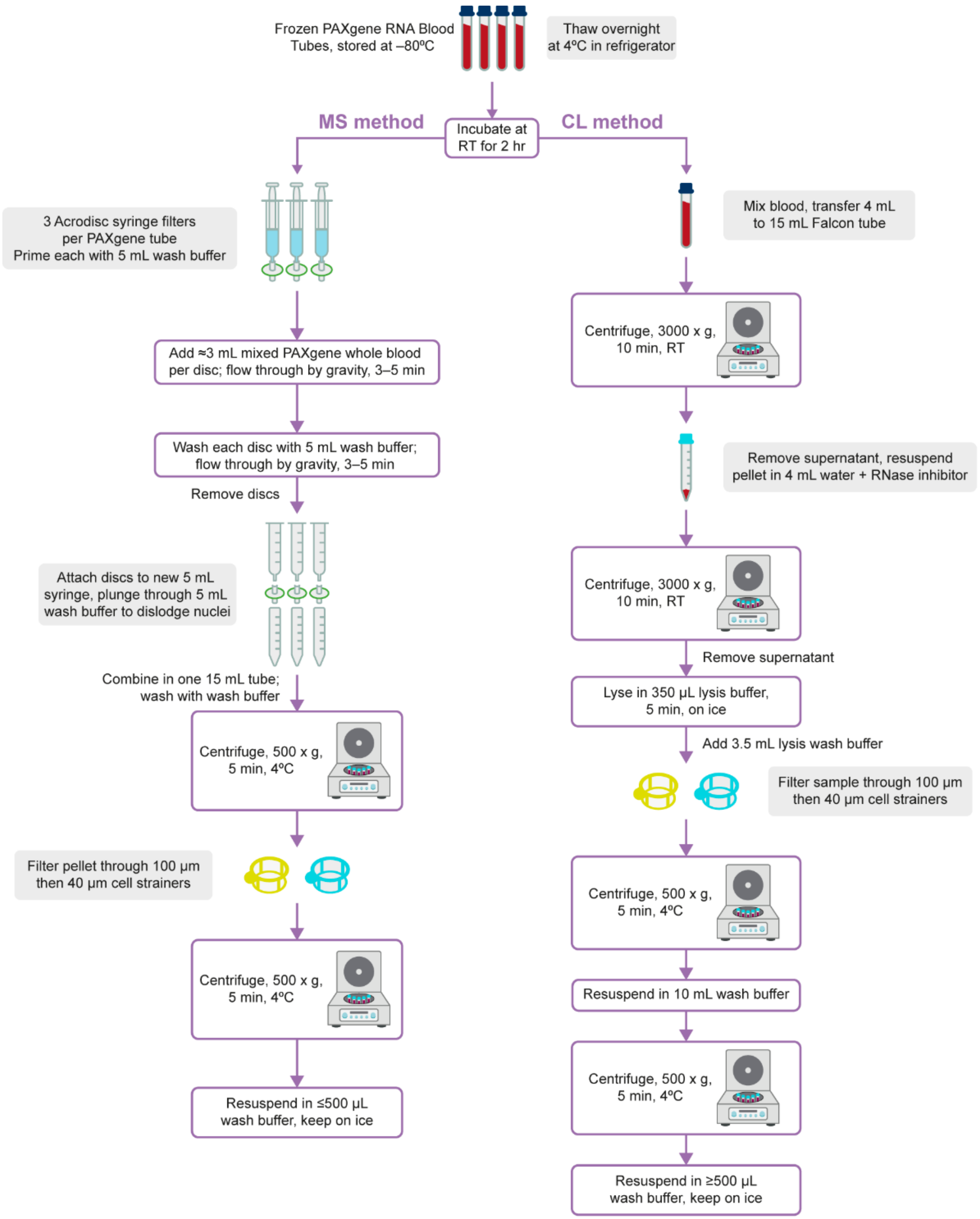
Frozen PAXgene RNA Blood Tubes, stored at –80ºC
Thaw overnight at 4ºC in refrigerator
Incubate at RT for 2 hr
MS method
CL method
3 Acrodisc syringe filters per PAXgene tube
Prime each with 5 mL wash buffer
Add ≈3 mL mixed PAXgene whole blood per disc; flow through by gravity, 3–5 min
Wash each disc with 5 mL wash buffer; flow through by gravity, 3–5 min
Remove discs
Attach discs to new 5 mL syringe, plunge through 5 mL wash buffer to dislodge nuclei
Combine in one 15 mL tube; wash with wash buffer
Centrifuge, 500 x g, 5 min, 4ºC
Filter pellet through 100 µm then 40 µm cell strainers
Centrifuge, 500 x g, 5 min, 4ºC
Resuspend in ≤500 µL wash buffer, keep on ice
Mix blood, transfer 4 mL to 15 mL Falcon tube
Centrifuge, 3000 x g, 10 min, RT
Remove supernatant, resuspend pellet in 4 mL water + RNase inhibitor
Centrifuge, 3000 x g, 10 min, RT
Remove supernatant
Lyse in 350 µL lysis buffer, 5 min, on ice
Add 3.5 mL lysis wash buffer
Filter sample through 100 µm then 40 µm cell strainers
Centrifuge, 500 x g, 5 min, 4ºC
Resuspend in 10 mL wash buffer
Centrifuge, 500 x g, 5 min, 4ºC
Resuspend in ≥500 µL wash buffer, keep on ice

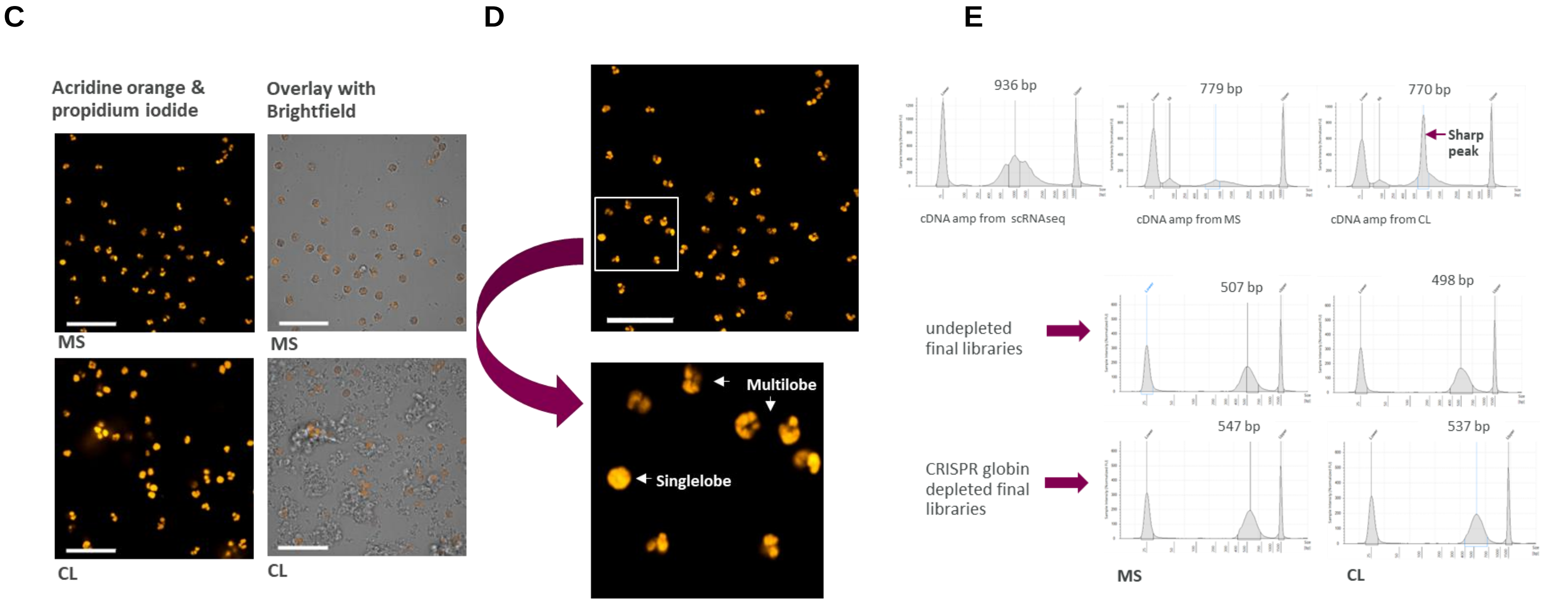


amp, amplified; bp, base pair; cDNA, complementary DNA; CL, cell lysis; MS, mechanical separation; RT, room temperature; scRNA-seq, single-cell RNA sequencing.

**Figure S2. Schematic of CRISPR depletion technology used to specifically digest globin genes (*HBA1*, *HBA2*, *HBB*, *HBD*)**

The method integrates in the 10x Genomics library construction workflow.

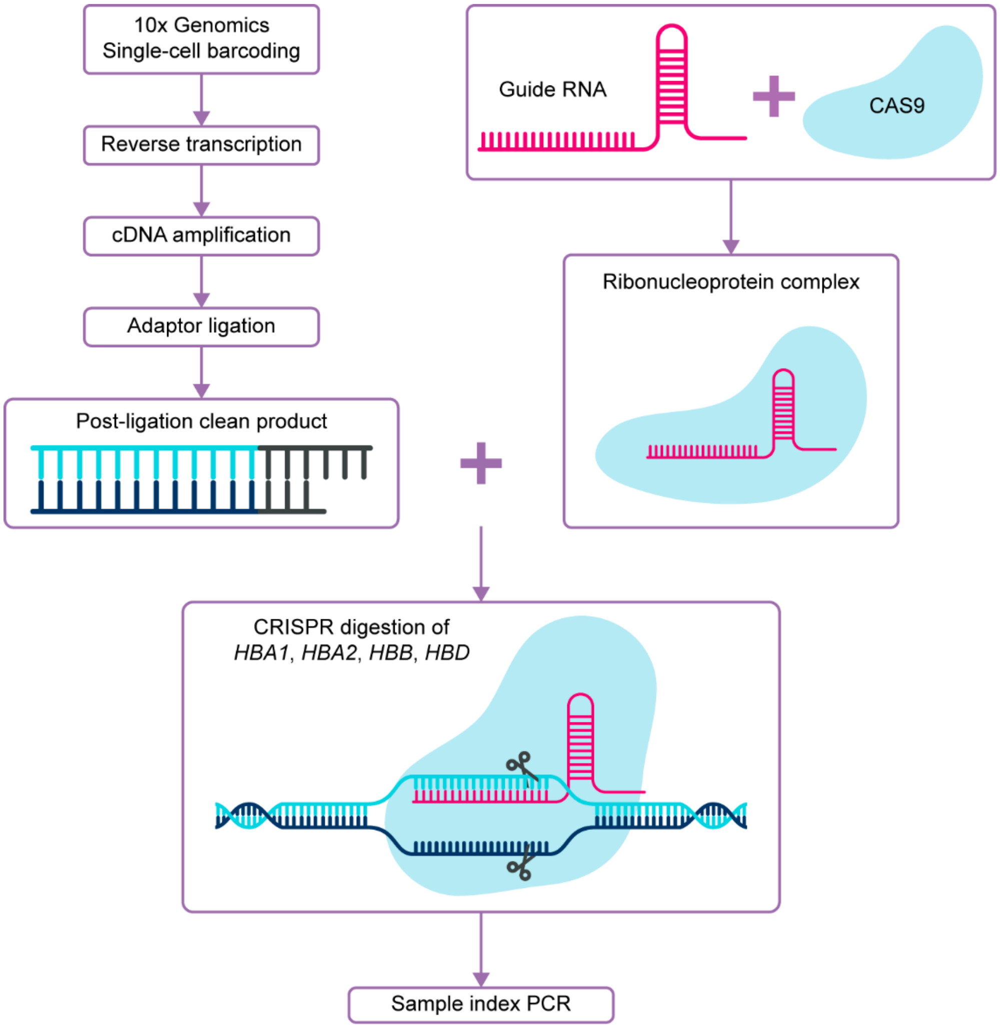


cDNA, complementary DNA.

### Figure S3. CRISPR-based globin transcript depletion across major cell types

Following nuclei separation by cell lysis (CL), reduction of globin transcripts by CRISPR (CL-D method) during final library construction resulted in substantial reductions in globin transcripts (dark mustard, CL-D, 2 samples per box) in B cells, CD4+ T cells, CD8+ T/NK cells, monocytes, neutrophils, and eosinophils compared with undepleted samples (blue, CL-undepleted, 2 samples per box). Horizontal line represents the median, box shows the interquartile range (IQR), and error bars represent 1.5 x IQR.

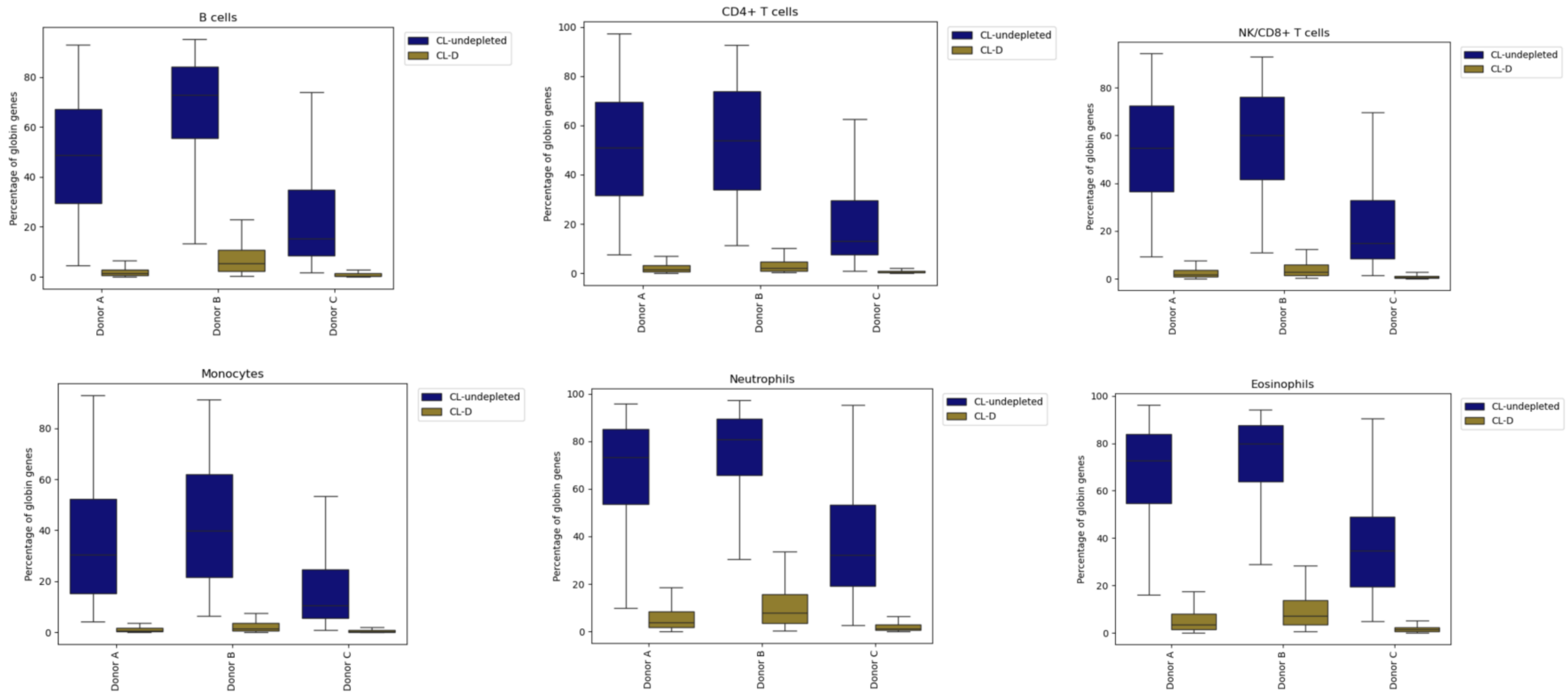
